# Causal attribution of agricultural expansion in a small island system using approximate Bayesian computation

**DOI:** 10.1101/2023.01.20.524853

**Authors:** Matt Clark, Jeffrey Andrews, Nicholas Kolarik, Mbarouk Mussa Omar, Vicken Hillis

## Abstract

The extent and arrangement of land cover types on our planet directly affects biodiversity, carbon storage, water quality, and many other critical social and ecological conditions at virtually all scales. Given the fundamental importance of land cover, a key mandate for land system scientists is to describe the mechanisms by which pertinent cover types spread and shrink. Identifying causal drivers of change is challenging however, because land systems, such as small-scale agricultural communities, do not lend themselves well to controlled experimentation for logistical and ethical reasons. Even natural experiments in these systems can produce only limited causal inference as they often contain unobserved confounding drivers of land cover change and complex feedbacks between drivers and outcomes. Land system scientists commonly grapple with this complexity by using computer simulations to explicitly delineate hypothesized causal pathways that could have resulted in observed land cover change. Yet, land system science lacks a systematic method for comparing multiple hypothesized pathways and quantifying the probability that a given simulated causal process was in fact responsible for the patterns observed. Here we use a case study of agricultural expansion in Pemba, Tanzania to demonstrate how approximate Bayesian computation (ABC) provides a straightforward solution to this methodological gap. Specifically, we pair an individual-based simulation of land cover change in Pemba with ABC to probabilistically estimate the likelihood that observed deforestation from 2018 to 2021 was driven by soil degradation rather than external market forces. Using this approach, we can show not only how well a specific hypothesized mechanism fits with empirical data on land cover change, but we can also quantify the range of other mechanisms that could have reasonably produced the same outcome (i.e. equifinality). While ABC was developed for use in population genetics, we argue that it is particularly promising as a tool for causal inference for land system science given the wealth of data available in the satellite record. Thus, this paper demonstrates a robust process for identifying the emergent landscape-level signatures of complex social-ecological mechanisms.

## 1 Introduction

The mosaic of land cover on the surface of our planet is a product of complex social-ecological dynamics that make up the complete land system (B. L. Turner, Lambin, and Reenberg 2007; B. L. Turner, Lambin, and Verburg 2021). Changes in Earth’s terrestrial surface have profound implications for ecosystem functioning and human wellbeing, and as such, are of critical importance to understand and predict (Steffen et al. 2006). A key challenge in understanding land cover change and designing effective policies is that there are often multiple plausible, and even interacting mechanisms that can cause a switch in land cover from one state to another (Lambin and Meyfroidt 2010). For example, in the case of agricultural frontier expansion, depletion of soil fertility in existing plots often promotes the conversion of nearby natural vegetation to new cropland, thus pushing the frontier outward (Casetti and Gauthier 1977). Agricultural frontier expansion into forested lands is also observed however, as a result of increased market value for crops and increased population pressure (P. Meyfroidt et al. 2018).

When designing policies to limit the conversion of natural areas into more intensive land use types such as rotational agriculture, it is important to determine the drivers of conversion, because the impact of a given policy will depend on the dominant driver. The introduction of new agricultural technologies, for instance, can have drastically different effects on the landscape depending on the primary driver of frontier expansion in a given system (see Kaimowitz et al. 1998 for a foundational review). If loss of soil fertility is the primary driver, new technologies can limit forest conversion as they allow existing plots to be farmed for longer periods of time, minimizing the need for agricultural operations to change location. However, if market forces are the primary driver of frontier expansion, this same intervention is likely to incentivize further forest conversion by increasing the returns from any given agricultural plot (P. Meyfroidt et al. 2018). Despite the importance of identifying causal processes in land cover change, actually doing so in any particular case has often proven difficult given the inherent social-ecological complexity of land systems (Patrick Meyfroidt 2016; B. Turner et al. 2020).

Land systems are complex adaptive systems characterized by feedbacks between the human and ecological subsystems, where a change in the state of one affects ongoing processes in the other and vice versa (Berkes, Folke, and Colding 2000; Le, Seidl, and Scholz 2012; Folke 2007). The possible distribution of land cover types in any one area is also highly path dependent, or constrained by past states and trajectories, sometimes further muddying the relationship between actual drivers and outcomes (Liu et al. 2007). Standard statistical tests fail to produce reliable inference in complex systems exhibiting feedbacks and path dependence, given that it is generally impossible to specify likelihood functions for such processes (Levin et al. 2013). Thus, while critical, identifying causal processes in land systems and social-ecological systems generally has proven difficult.

To begin to build causal theory in complex land systems, researchers commonly use computer simulations to abstract key phenomena and produce ‘what-if’ scenarios (Ahimbisibwe et al. 2021; An et al. 2021). Simulations allow researchers to code complexities such as feedbacks and path dependence directly into a model of the processes under examination, in order to formally define a hypothesized causal mechanism and check the logical implications and internal validity of their assumptions (Epstein 2008; Verburg 2006). While simulations like this are important for theorizing about social-ecological systems, it can be difficult to relate them back to empirical data and tell where exactly the real-world sits in the multidimensional parameter space of the model (Ren et al. 2019). Without this information, we are limited in our knowledge of how well a given simulation accurately distills the processes we are hoping to examine, and how we might use such a model to infer important things about the real-world.

The biological sciences have largely led the development of methods for comparing simulated, theoretical causal processes to observed data. Ecological research in particular has made considerable use of simulation modeling to theorize about how complex interactions among individuals lead to observed patterns at the population level (i.e. individual-based modeling) (Grimm 1999; DeAngelis and Grimm 2014; Grimm and Railsback 2013). Relatively early on in the use of individual-based modeling in ecology, researchers developed the general process of pattern-oriented modeling in which a given hypothesis is evaluated on its ability to recreate an observed biological pattern at an appropriate temporal and spatial scale (Wiegand et al. 2003; Grimm et al. 2005). This method allowed researchers to match observed trends in population change with plausible rates for various demographic parameters such as pre-breeding survival in woodpeckers, road mortality in lynx, and annual male survival in amphibians, among many others Wiegand et al. (2003). While pattern-oriented modeling provides a general structure for interrogating causal hypotheses of complex phenomena with empirical data, it does not adequately account for stochasticity in the outcomes of hypothesized mechanisms. In particular, this method fails to quantify the complete range of model parameters that may reasonably reproduce observed patterns and the frequency in which they do so.

Toward this aim, approximate Bayesian computation (ABC) has emerged as a formal method of pattern-oriented modeling in which researchers run simulation models across many parameter values and systematically accept or reject the outputs of each run as consistent with observed data. All accepted parameter values are then aggregated into a probability distribution of parameter values that are likely to produce the observed data (Hartig et al. 2011; Troost et al. 2022). ABC has proven effective for identifying the range of simulation model parameters consistent with observed data across a wide breadth of biological fields from ecology to epidemiology (Scranton, Knape, and Valpine 2014; Kosmala et al. 2016; Vaart, Johnston, and Sibly 2016; Boult et al. 2018; Martínez et al. 2011; Cipriotti et al. 2012). When different simulation parameters represent specific hypothesized causal processes, ABC can then be used to estimate the probability that a given causal process produced a given set of observed data. Importantly, ABC enables researchers to statistically estimate model parameters even for generative models containing complex processes such as the feedbacks and path dependence characteristic of social-ecological systems (Gallagher et al. 2021).

In this paper we demonstrate the utility of ABC for generative inference in complex land systems and social-ecological systems generally. Specifically, we simulate hypothesized patterns of agricultural expansion in a small island system given two possible drivers, declining soil fertility and external market forces. We then filter the range of possible model parameters to just the inputs that produce land use patterns consistent with the observed time-series of agricultural frontier expansion. We show that this method allows us to determine the proportion of each of these drivers in causing the observed agricultural expansion in our study system, Pemba Island, Tanzania. In this way, we provide a straightforward demonstration for linking land system simulations with empirical data to draw causal inference in even very complex systems involving feedbacks and path dependence.

## 2 Study system

### 2.1 Pemba

The Indian Ocean archipelago of Zanzibar is a semi-autonomous jurisdiction lying off the coast of Tanzania. Pemba, the northernmost island, is densely populated with 428 people per square kilometer. While the island has a few main population centers (Wete, Chake Chake, and Mkoani), the vast majority of the island’s 400,000 people live in small villages scattered across some 120 wards (*shehia*), all of which are connected by a dense, relatively modern road network (fig. 1). Based on the 2022 census, we estimate the average growth rate between 2012 and 2022 to be about 2.1%, more than double the world average of 0.9% per year.

**Figure 1:**
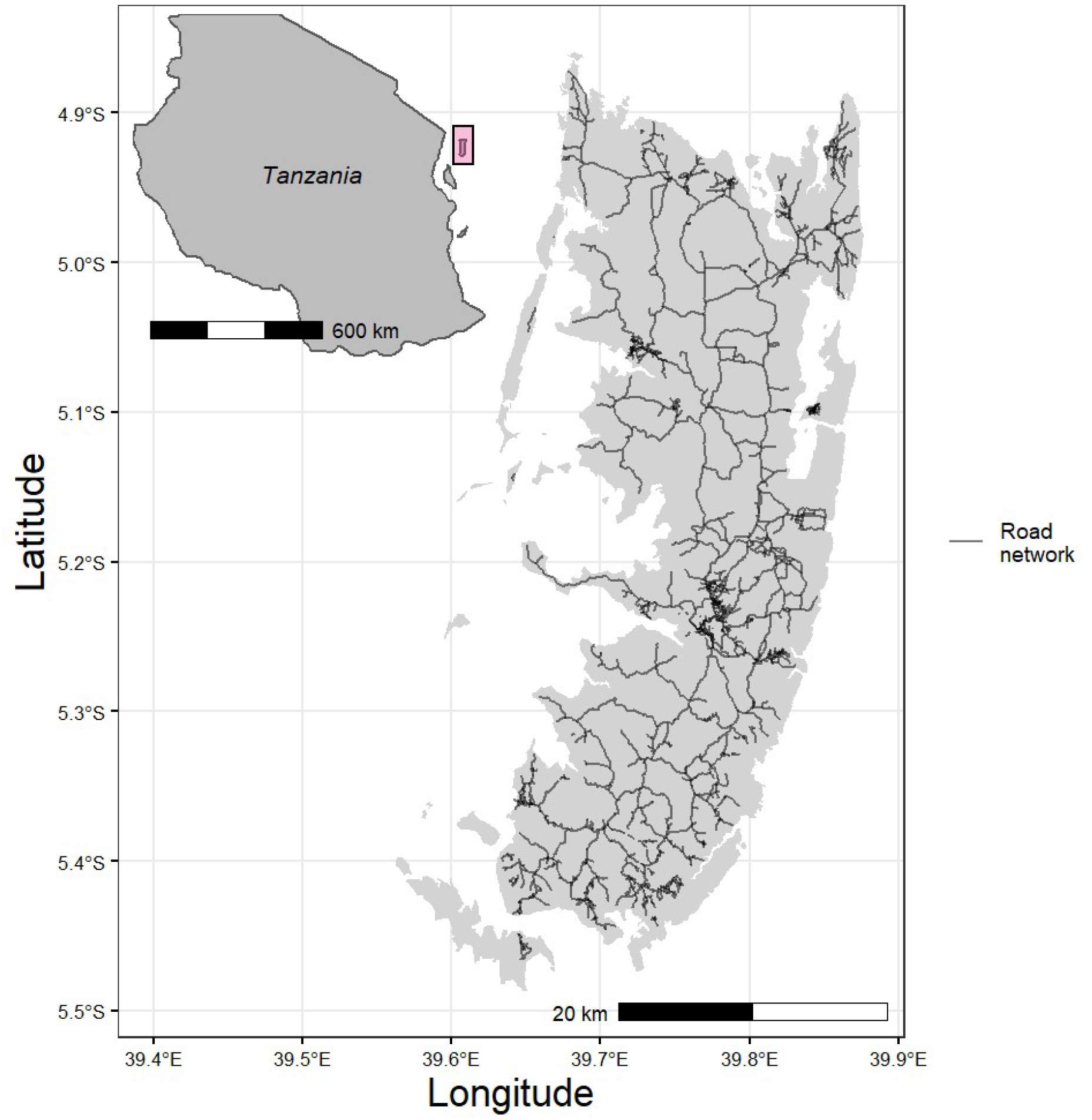
Inset map showing the location of Pemba island relative to the Tanzanian mainland (upper left) and the dense Pemban road network (main figure).

The rural economy of Pemba is primarily subsistence (farming, fishing, and livestock), with a single important cash crop, cloves. Cloves were introduced to the island in the early 19th century and have come to dominate the economy of the island’s western half due to the region’s highly productive soils (see below). In contrast, the island’s eastern side, characterized by poor, shallow soil, has lower human population density, is less developed, and is generally less economically productive. For example, in an extensive household survey focused on economic production carried out in 2017, we find that the average household income from the sale of crops (excluding cloves) in the east is approximately $25, compared with an estimated $80 in the highly productive western half.

Nevertheless, there is considerable pressure in the east for new farmland. Some 30% of All eastern households surveyed stated that they had cleared forested land to expand their farming operations in the past 7 years. On average, households report clearing approximately 0.91 acres, primarily to plant staple crops such as cassava. And while cassava is almost exclusively a subsistence crop in Pemba, there has been considerable pressure to develop the Pemban economy in the past decade, as it has lagged behind the rest of Tanzania and Zanzibar. New development initiatives, particularly in agriculture, are a constant of government programming, and new crops such as watermelon and tomatoes are being experimented with on the once underutilized eastern soils. However, Pemba has historically struggled to develop its own internal market for agricultural goods, and the impact that these new cash crops are having on the eastern landscape is unknown.

### 2.2 Coral rag vegetation and rotational agriculture

Pemba is a narrow island, in many places just 15 kilometers wide, yet most of the environmental variation exists across the narrow east/west span. This is owed to three distinct soil types that run the length of the island and can generally be thought of as going from deep and fertile in the west, to shallow and nutrient poor in the east (Stockley 1928). The easternmost topography is characterized by jagged, fossilized coral beds covered with a shallow soil layer and scrubby vegetation ranging from approximately 1 to 5 meters in height (i.e. coral rag forest) (Burgess and Clarke 2000; Burrows et al. 2018). This forest type has traditionally been overlooked by conservation efforts in Pemba, and Zanzibar generally, yet it is critical habitat for a variety of plant and animal species such as the endemic Pemba flying fox (*Pteropus voeltzkowi*) (Kingdon 1988).

The shallow soils atop porous coral rag geology characteristic to eastern Pemba cannot be productively farmed for long, continuous periods. Farmers in this region thus typically rely on rotational swidden agriculture where primary forest is cleared and the land is farmed for a short period, then left to recharge for a number of years (fig 2). While this process could potentially be interrupted by regenerative agricultural practices or the introduction of rainwater catchment systems, currently farmers in this region lack resources to escape the cycle of ecologically costly, short-term resource extraction (Wild et al. 2020; Biazin et al. 2012; Garrity et al. 2010). Thus, in this system, there are considerable feedbacks between the condition of the environment, decisions made by farmers to clear and farm a forested area, and the state of the environment in future time periods. Further, as any one forested area is cleared, it opens new patches to potential clearing through frontier expansion. Hence, specific patches available in any one time period are highly dependent on social-ecological events in previous time steps. Thus, this system displays both the feedbacks and path dependence characteristic of complex social-ecological systems.

**Figure 2:**
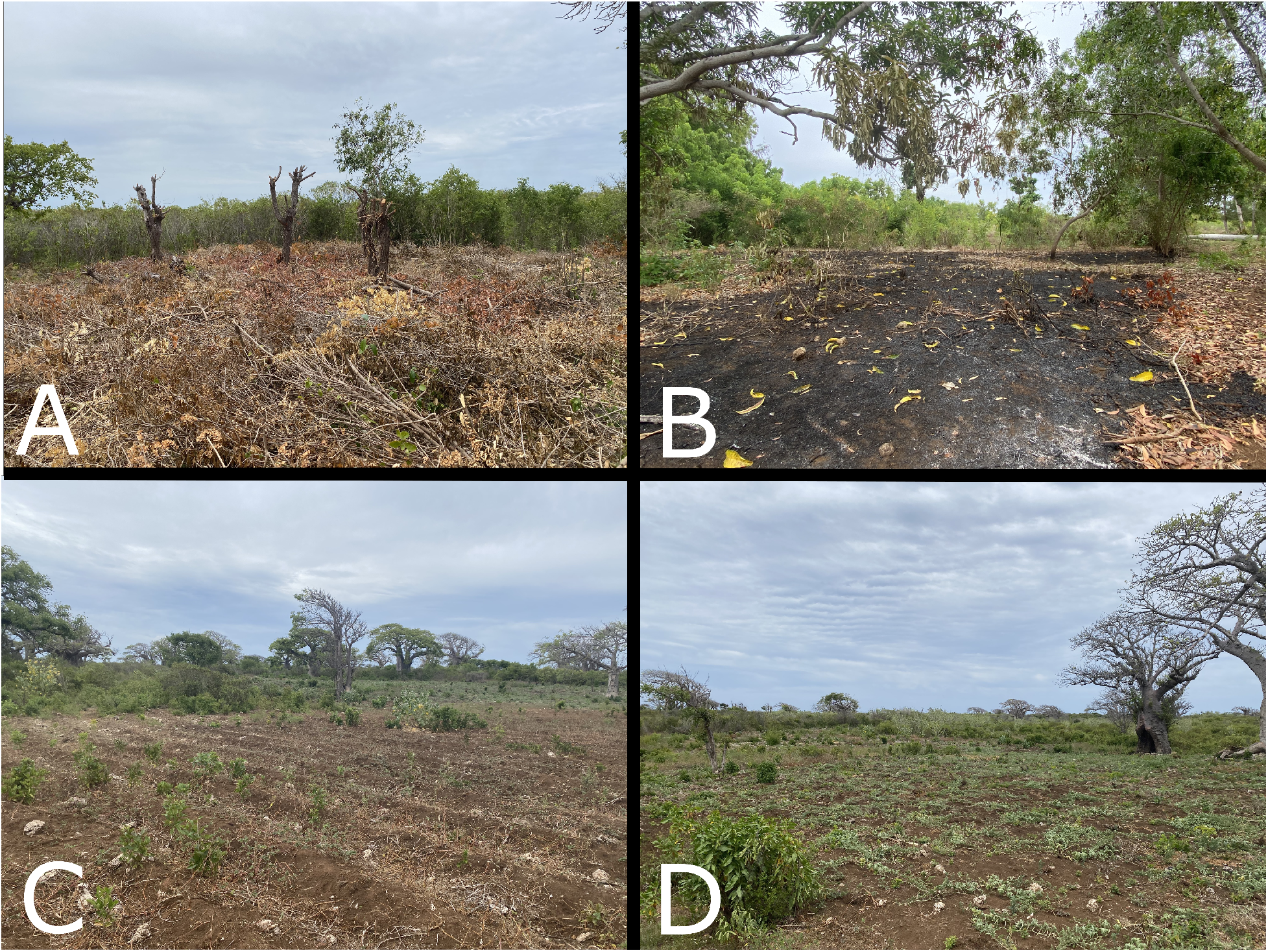
Panels A, B, C, and D show the process of coral rag forest conversion to agriculture in Pemba in four distinct stages. Panel A shows the cutting of coral rag vegetation, which is then left to dry before burning. Panel B shows a freshly burned plot before planting. Panel C shows a productive agricultural field. Panel D shows a fallow agricultural field.

## 3 Methods

### 3.1 Data

#### 3.1.1 Land cover classification

We produced 20 m land cover maps using top of atmosphere Sentinel-2 time series imagery and ancillary datasets in a data fusion approach using Google Earth Engine (GEE) (Gorelick et al. 2017; Mondal et al. 2019). GEE is an open-access cloud computing platform that hosts petabytes of freely available earth observation data, and is ideal for creating land cover maps with built-in classification functions. For this study, we created annual median composite images for conversion into thematic land cover maps. We filtered the time series by date, and used the image metadata to further filter by estimated cloud cover, using a threshold of <20%. We then used the ‘QA60’ quality band to remove any remaining clouds prior to creating the composites of median values. Beyond the native Sentinel-2 spectral bands, we calculated common normalized difference indices useful for distinguishing common land cover types such as the Normalized Difference Vegetation Index (and red edge adaptations) and the Normalized Difference Water Index (NDWI), as well as its modified version (MNDWI) (DeFries and Townshend 1994; Schuster, Förster, and Kleinschmit 2012; Xu 2006; Gao 1996). We also used synthetic aperture radar (SAR) backscatter from the corresponding Sentinel-1 ground range detected time series available in GEE. We used SAR scenes from ascending paths only and incorporated both vertical-vertical and vertical-horizontal polarizations in our analysis. SAR data are known to be influenced by varying incidence angles, so we normalized these images by multiplying the backscatter by the incidence angle with the understanding that greater incidence angles result in less backscatter returned to the instrument (Banks et al. 2019; Kaplan et al. 2021). To reduce inherent speckle in the SAR images, we opted for a time for space substitution by using a mean composite of all images for the given year to maintain a 10 m spatial resolution. Lastly, we considered topographic covariates for classification (elevation, slope, and aspect derived from the NASA Shuttle Radar Topography Mission digital elevation model) that dictate locations of land covers of interest relative to sea level and topography (Table S1) (Farr et al. 2007).

From each composite image, we collected representative training samples for all classes of interest (mangrove, high forest, agriculture, urban, bare, coral rag, other woody vegetation, and water). We trained a random forest classifier using training samples from 2018, 2019, & 2021 to account for potentially varying atmospheric and illumination conditions among images (Breiman 2001). The random forest we used for classification had 100 trees, and utilized four variables per split (the square root of the number of covariates), consistent with other remote sensing applications (Belgiu and Drăgut 2016). Due to relative class imbalances, we chose to use a stratified random sampling design to assess the accuracy of our outputs, and computed area adjusted accuracy metrics (Olofsson et al. 2013; Stehman and Foody 2019). An expert hand labeled these stratified points based on high resolution median composite PlanetScope images with 20 points from each mapped class. Results show estimated overall accuracies of 92.86% (+/-4.21%) and 95.93% (+/- 2.95%) for 2018 and 2021, respectively (Table S2, Table S3). Much of the confusion and sources of error in the maps is found among upland woody vegetation classes (high forest, mangrove, and other woody vegetation) that share similar spectral and physical characteristics. Other mentionable errors occur among urban and agriculture in 2018, and bare and agriculture in 2021, leading to relatively high uncertainty for area estimates and producer’s accuracies in the rare urban and bare classes in 2018 and 2021, respectively (Table S4, Table S5).

#### 3.1.1 Interview data

In July 2021, we conducted informal interviews with staff of Community Forests Pemba, a nonprofit aimed at building conservation capacity on the island, and farmers in four *shehia* with coral rag forests in the east of the island. Researchers asked farmers about how they make decisions regarding when and how long to farm and fallow agricultural plots, as well as how they decide to clear forest vegetation to establish new cropland. There was broad consensus among non-profit staff and farmers that agricultural plots in these areas are typically farmed for two years and then left fallow for three years. Clearing forested land is labor intensive and thus, forested land is only cleared when nearby productive plots reach their two-year limit and no other previously-cleared adjacent plots are available.

### 3.2 Individual-based simulation

Our individual-based model incorporates two mechanisms of land conversion from coral rag vegetation to productive agricultural land. The first follows the decision rules described during the farmer interviews where coral rag vegetation is converted in response to soil degradation and space limitations following the fallowing of cropland. Under this mechanism, each pixel (20m area) is autonomous and follows the following basic set of decision rules also described visually in figure 4. Each pixel classified as agriculture in the study *shehia* is initialized randomly as either productive or fallow. Each productive agricultural pixel is then randomly assigned to either the first or second year of agricultural production. Each fallow agricultural pixel is randomly assigned as in the first, second, or third year of fallow time. When transitioning to the next year, agricultural pixels in their second year of production go fallow and the lost agricultural production is relocated as follows. If there is an adjacent (8 directional) fallow pixel in the final (third) year, the crop production moves there. If there is no adjacent third year fallow pixel available and there is adjacent coral rag vegetation, that coral rag pixel is converted to first year productive agriculture for the next year. All other land cover types (e.g. water, urban, mangrove, etc.) are left alone.

**Figure 3:**
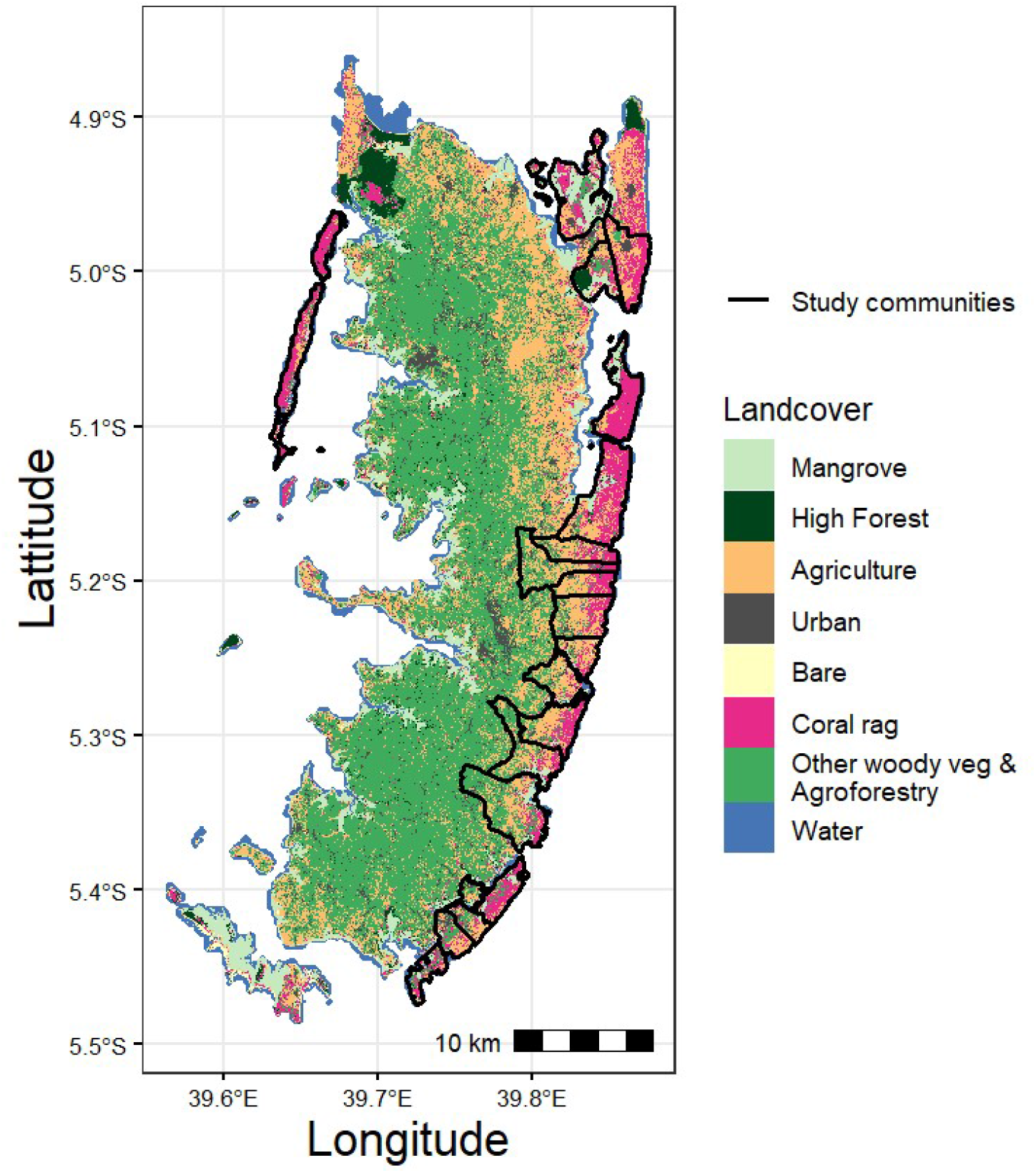
Land cover estimates for the year 2018 in Pemba. The classification for each 20m pixel is distinguished by color, with the class of interest, coral rag vegetation, highlighted in pink. Black lines show each of the 19 *shehia* included in this study.

**Figure 4:**
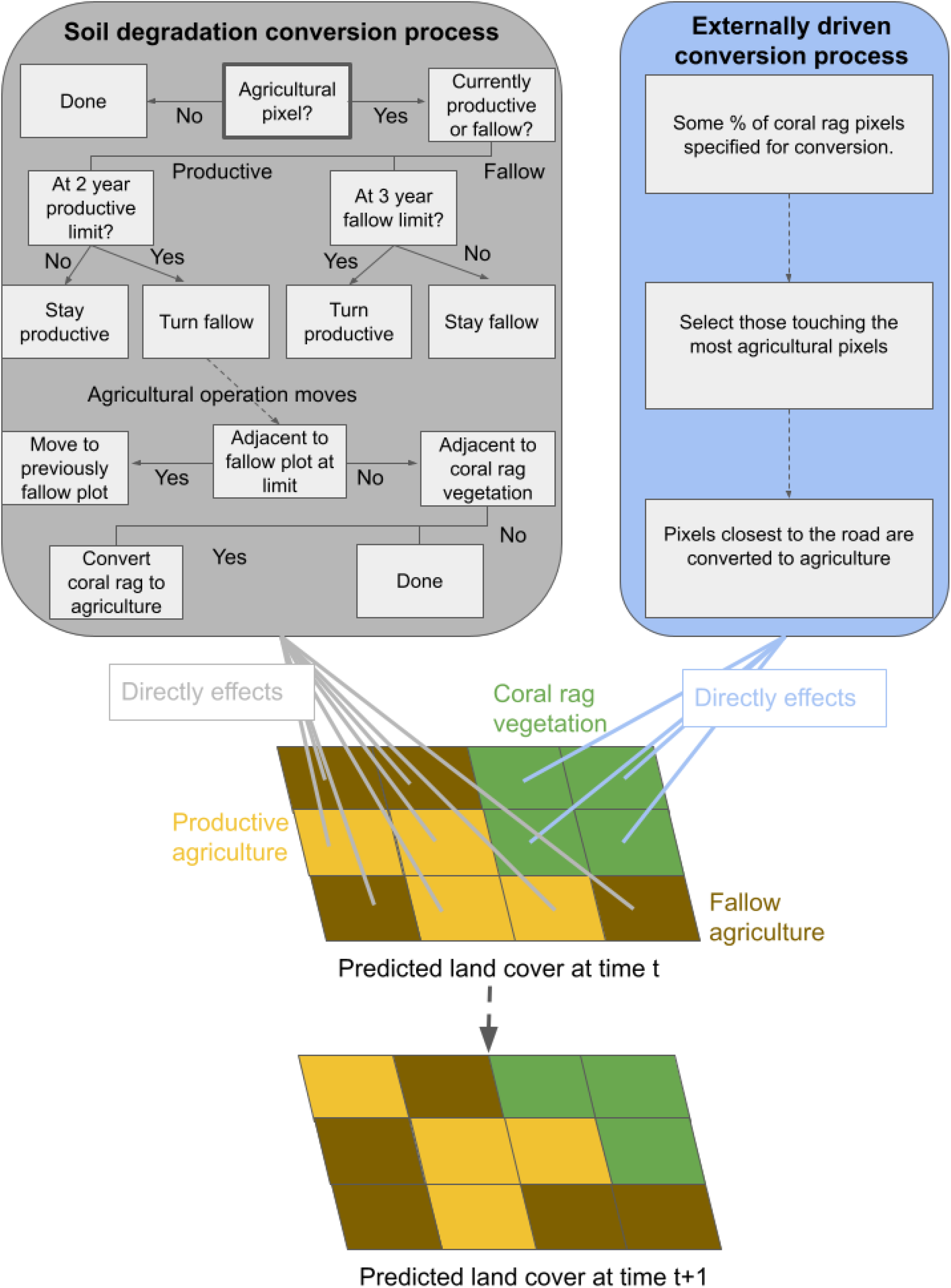
Visual representation of the individual-based simulation used to model coral rag vegetation conversion to agriculture in each of the 19 *shehia* included in this study. The top two boxes show the competing causal forces driving land conversion. The bottom portion of the figure shows how those forces affect the pixel-based land cover in each time step.

The second mechanism of land conversion from coral rag vegetation to productive agricultural land represents all other factors driving land conversion outside of the soil degradation process. In our system, the most prominent other factors include a rapidly growing population and the increasing market value for crops. Under this mechanism, some additional percentage of coral rag vegetation is converted to first year agriculture each year, representing a yearly rate of forest loss caused by factors other than soil degradation. Coral rag pixels allocated to conversion are those that are adjacent to the greatest number of agricultural pixels. When coral rag pixels are adjacent to an equal number of agricultural pixels, then the coral rag pixels closest to a road are selected for conversion to first year agricultural land. A visual representation of this simulation can be found in figure 4.

For each run of this simulation, the observed land cover on Pemba Island in 2018 (fig. 3) is used as the starting year and the model is run for three time steps to yield a predicted land cover for 2021, given a set rate of externally driven agricultural expansion. By then comparing this predicted land cover in 2021 to the observed land cover in 2018, we can produce a predicted number of coral rag pixels to be converted to agriculture under different rates of externally driven land conversion.

### 3.3 Approximate Bayesian computation (ABC)

As described in the introduction, simulation modeling allows researchers to formally express complex, hypothesized causal mechanisms in land system science. The primary limitation for the use of simulation modeling to enhance our understanding of real-world causality, however, is the absence of a straightforward statistical process for relating simulations to empirical data. Approximate Bayesian computation is one way to produce such generative inference (Kandler and Powell 2018). In this framework, researchers run a simulation model under a wide range of parameter combinations, representing alternative hypotheses, to produce many simulated datasets for which all parameters and outcomes of interest are known. All simulated datasets are then systematically accepted or rejected as consistent with the observed data, and the combination of model parameters that provides the best fit is estimated probabilistically (Vaart et al. 2015; Beaumont, Zhang, and Balding 2002). Hence, ABC allows researchers to quantitatively compare the likelihoods of competing hypothesized causal mechanisms in complex land systems.

In this study we vary just one model parameter, the externally driven rate of coral rag vegetation conversion to rotational agriculture. We first specify a prior distribution that we believe will capture all possible values of this parameter of interest (fig. 5). This prior is based on a combination of calibration with earlier models, and *a priori* understanding of the system from working with local conservation organizations and farmers.

**Figure 5:**
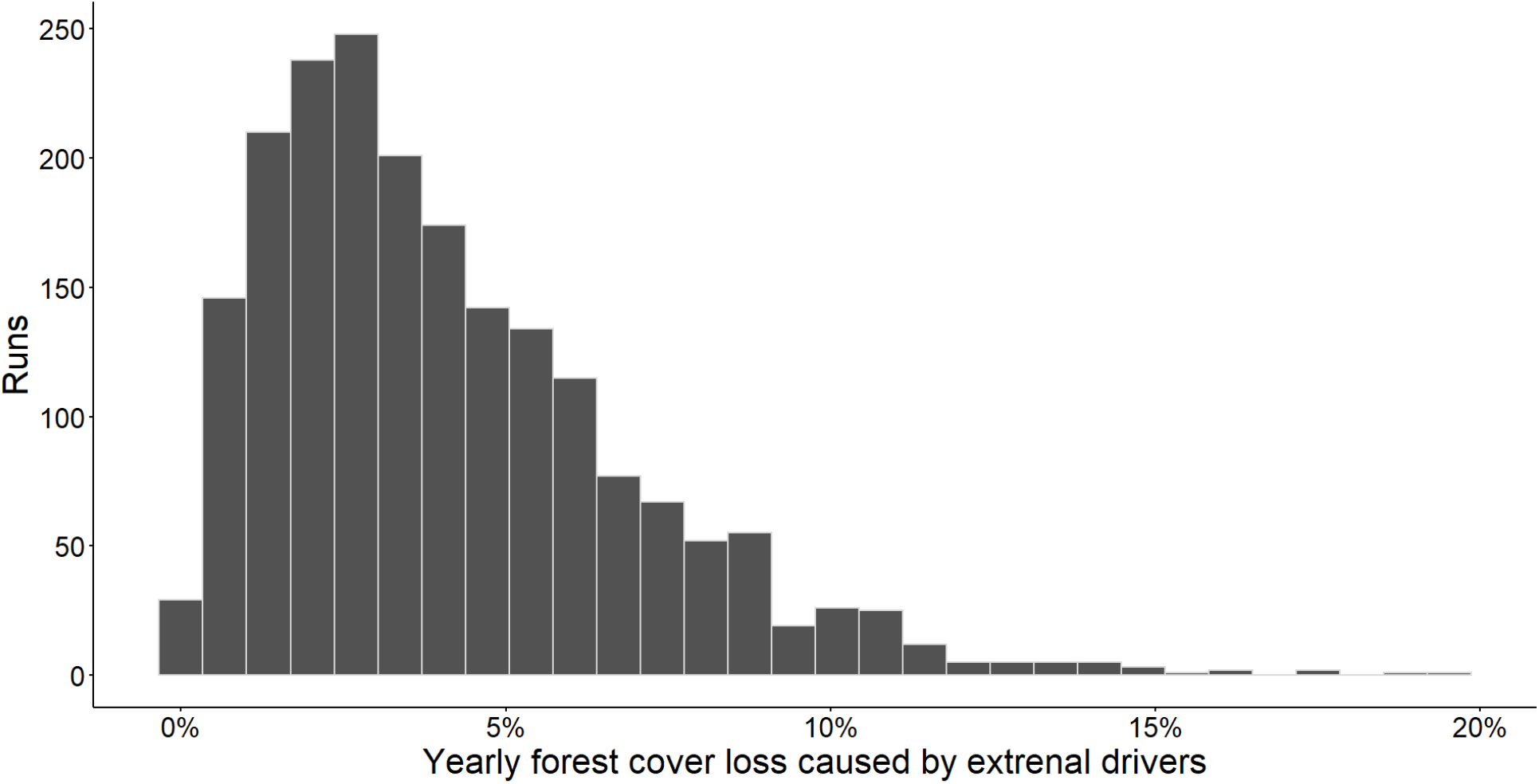
Two thousand draws from the prior distribution of the rate of externally driven coral rag vegetation conversion to agriculture. These draws were used to produce the prior distribution of the expected number of coral rag vegetation pixels to be converted to agriculture from 2018 - 2021 for each *shehia*.

We then run our simulation model 2,000 times using draws from this prior distribution as the parameter of interest — the externally driven rate of coral rag forest conversion. For each run, the model then produces a synthetic dataset including the number of expected agricultural conversions for each *shehia* in our study. Each of these synthetic datasets is then accepted or rejected as consistent with the observed changes in land cover as measured through our 20m land cover classification map for 2021. We use an acceptance criteria of predicted agricultural conversions within 10% of the observed conversions. Finally, the parameter value for the extrinsic growth rate in each synthetic dataset that is accepted as consistent with the observed data is saved as one “draw” from the posterior for the estimated real-world parameter value.

## 4 Results

### 4.1 Parameter estimation

As described in the methods, for each *shehia* we ran 2,000 simulations, using each draw from the prior distribution of externally driven coral rag forest conversion for each simulation (figure 5). Each of these simulations converted some number of coral rag vegetation pixels to agriculture in each of the 19 *shehia* from 2018 to 2021 (fig. 6). An average of 231 of these estimates per *shehia* were within 10% of the observed number of converted coral rag vegetation pixels (Sunnåker et al. 2013). Keeping only these synthetic datasets consistent within the 10% error bound, we observe the distribution of externally driven growth parameter values that, based on our model, are likely to have produced the observed 2021 land cover (fig. 7). The median parameter values consistent with the observed data (parameter estimates) for each *shehia* range from 0.0% to 3.9% for yearly coral rag forest cover loss due to external forces (fig. 7). For four of the study *shehia*, the observed number of coral rag forest pixels converted to agriculture was fewer than predicted by the cycles of soil degradation built into our model alone. These four *shehia* all generally exhibited overall low rates of observed coral rag forest conversion to agriculture observed through the satellite imagery. We discuss contextual factors that may be influencing these trends in the discussion.

**Figure 6:**
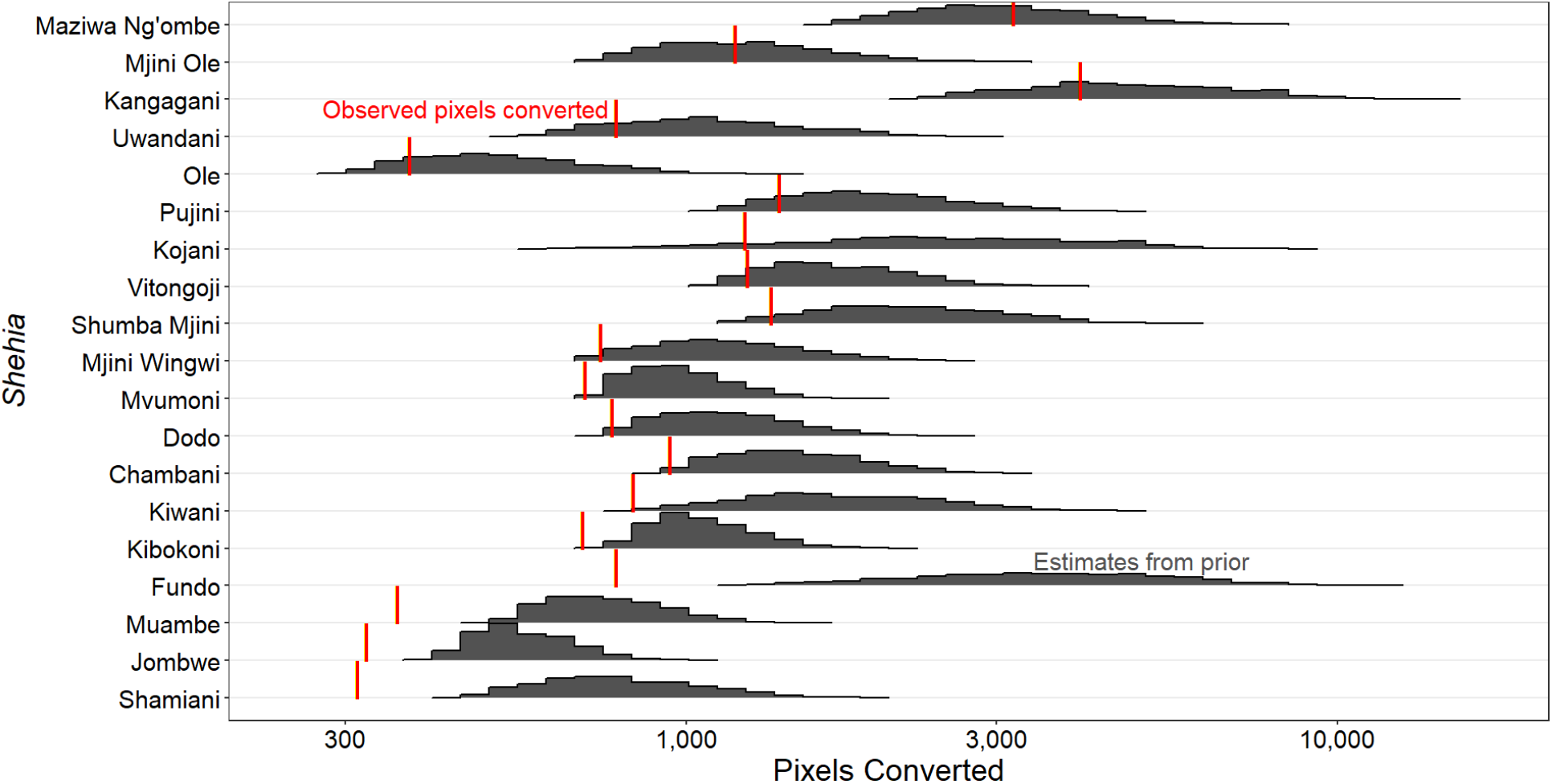
Black histograms represent the expected number of coral rag vegetation pixels to be converted to agriculture for each draw from the prior for each *shehia* from 2018 to 2021. Red lines show the observed number of conversions for each *shehia* from 2018 to 2021.

**Figure 7:**
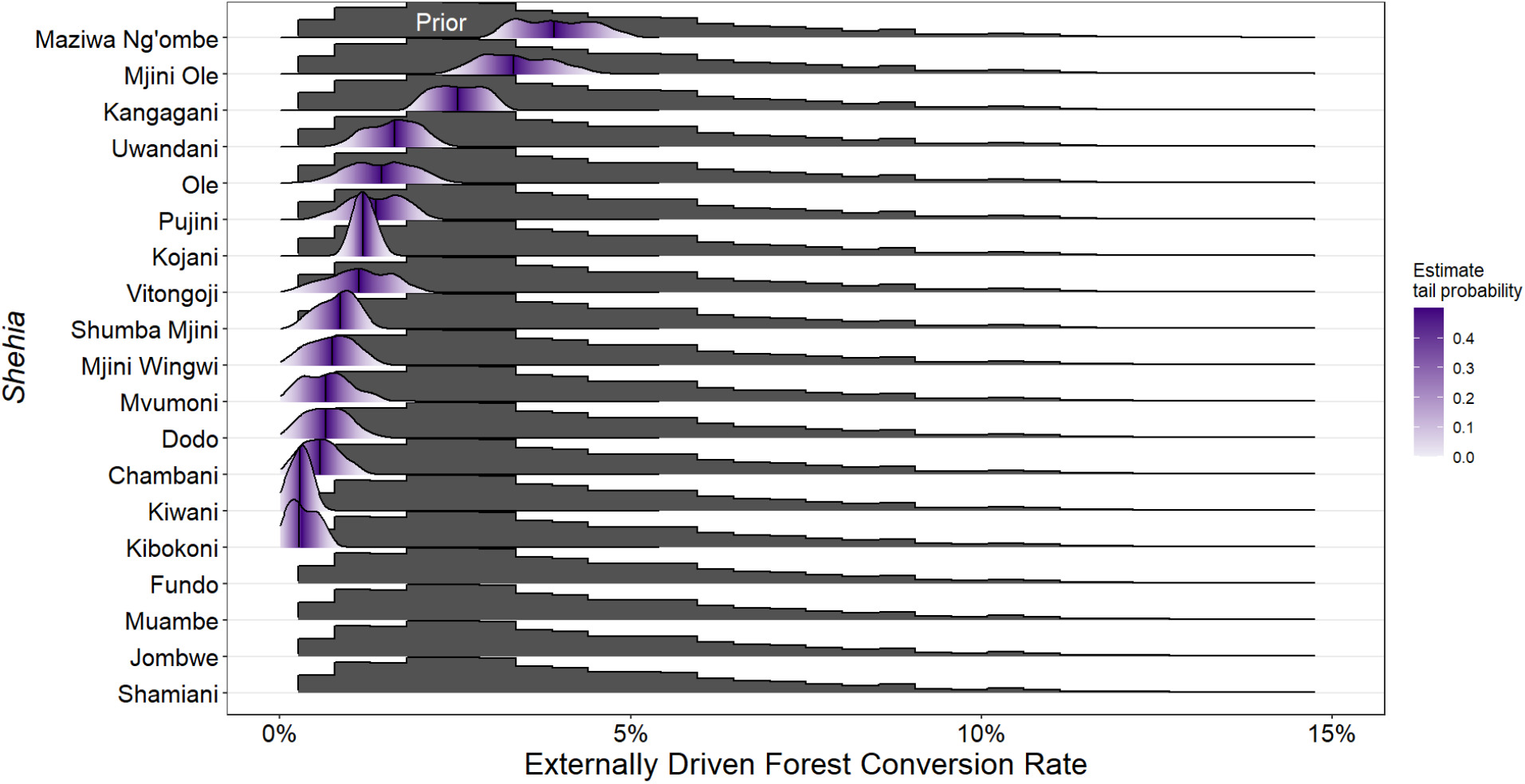
Black histograms show the prior distribution of externally driven deforestation rates for coral rag vegetation in Pemba. The prior values are identical for each *shehia*. Purple density plots show the density of draws from the prior in which the prior value resulted in a deforestation rate within 10% of the observed rate for that *shehia*. Shading in the density plots shows the tail probability that a particular value from the prior will result in the observed land cover based on our model.

The width of the posterior parameter estimates shown in figure 7 are indicative of how much information about causal processes we can infer from the observed land cover change from 2018 to 2021. Wider estimates indicate a greater degree of equifinality, where a wide range of externally driven deforestation rates could have produced the observed data. Conversely, narrow parameter estimates indicate that only a small range of externally driven deforestation rates could have produced the observed number of coral rag vegetation pixels converted to agriculture from 2018 to 2021. Thus, when parameter estimates are more narrow, the data carry a stronger underlying causal signature. The width of the posterior parameter estimates can then be thought of as the range of processes that could have reasonably produced the observed data (Kandler and Powell 2018). Across *shehia*, we observe relatively narrow parameter estimates, with an average range of 1.47%; the most narrow being 0.46% and the widest being 2.24% (fig. 7). We can conclude then that based on our model, on average, less than a 1% increase or decrease in the externally driven deforestation rate from the median model estimate is likely is result in the observed land cover change in each *shehia*.

### 4.2 Estimating the contribution of each process

With the observed rates of coral rag vegetation conversion from 2018 to 2021 and estimates for the externally driven agricultural expansion (deforestation) rate for each *shehia* in hand, we can assess the proportion of total loss driven by soil degradation versus external influences such as market integration and population growth. We subtract the median estimate of the contribution of external forces from the total satellite-observed rates of conversion to yield the point estimates shown in figure 8.

**Figure 8:**
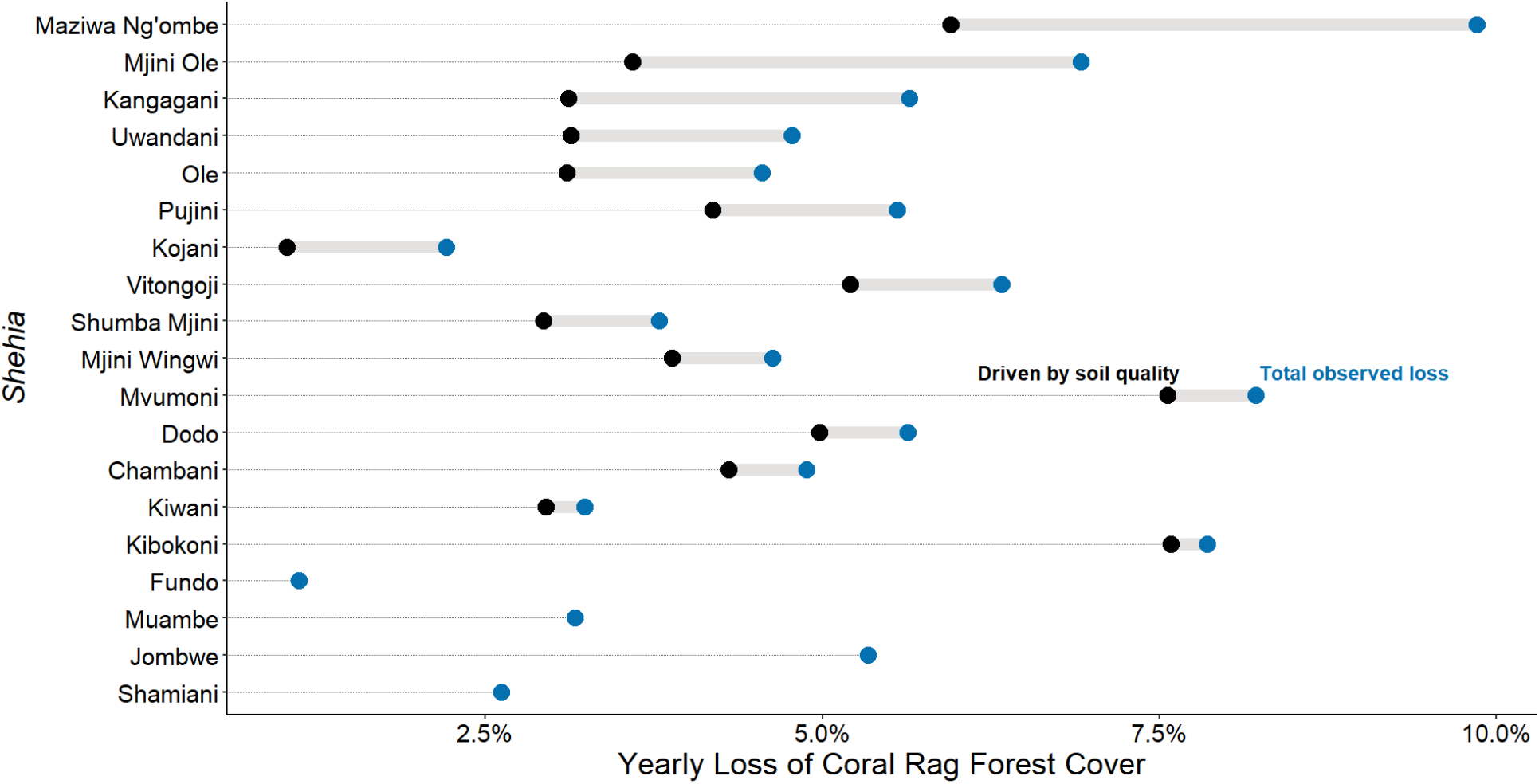
Barbell plot showing the median estimates of the influence of soil degradation on the observed coral rag vegetation loss in each *shehia*. Blue points show the total loss of coral rag vegetation as measured using satellite imagery. Black points show the median model estimates for the contribution of soil fertility loss to the observed deforestation.

On average, we observe a 5.1% yearly rate of total coral rag forest conversion to agriculture from 2018 to 2021. Between study *shehia* this rate of total conversion ranges from 1.1% to 9.8%. We estimate that the percent of coral rag vegetation converted to agriculture in each *shehia* driven by soil degradation was between 1.0% and 7.6% per year, with an average rate of 4.0%. By comparison, a relatively small proportion of coral rag vegetation in each *shehia* is converted to agriculture each year as a result of external forces. We estimate that the average rate of externally driven conversion is 1.1% of total coral rag forest cover, ranging from 0.0% to 3.9% between *shehia*.

While there is considerable variability in both the total rate of coral rag vegetation loss and in the contribution from external drivers between *shehia*, a general and intuitive trend is that *shehia* with a greater proportion of loss caused by external drivers show greater coral rag vegetation loss overall. The four *shehia* that showed a total number of observed coral rag pixels lost below that which is expected from soil degradation alone in figure 5, also show relatively little vegetation loss as a percentage of total coral rag cover from 2018 to 2021.

## 5 Discussion

### 5.1 Pemba specific implications

Across the *shehia* in our study, we observe a 5.1% yearly rate of coral rag vegetation conversion to agriculture. We show here that reported farmer behavior in response to soil degradation, paired with the topography of the study *shehia*, should account for an estimated 4.0% yearly rate of conversion on average. Thus, while variable, we can conclude that on average a relatively small proportion of the observed coral rag vegetation conversion to agriculture in the study *shehia* is driven by external forces such as increasing demand driven by increasing subsistence needs or market forces. This finding matches our theoretical expectations given that farmers in this region generally clear land in order to plant low value staple crops such as cassava (P. Meyfroidt et al. 2018). This suggests that regenerative agriculture programs, along with rainwater catchment systems may considerably reduce the long-term loss of coral rag forest in Pemba, Tanzania. These programs will also ease the strain that clearing forested land puts on farmers, and may potentially help farmers break free of the cycle of environmentally damaging agricultural practices in pursuit of short term gains.

While the 1.1% average estimated yearly conversion rate of coral rag vegetation to agriculture in the study *shehia* driven by external forces is relatively small compared to the rate of loss driven by soil degradation, it is not negligible. Further, this value could reasonably increase as a result of continued market integration and population growth in Pemba. As the impact of external forces on land conversion increases, theory tells us that the effect of improved agricultural technologies on mitigating forest loss may be reduced or even reversed (Kaimowitz and Angelsen 1998). Top-down interventions such as designating one or more coral rag forest reserves on the island may help to slow the conversion of primary forest in some areas, but may also be subject to leakage, attenuating their overall efficacy (P. Meyfroidt et al. 2020; Bastos Lima, Persson, and Meyfroidt 2019). Instead, interventions focused on the introduction and establishment of value chains for alternative income sources, aside from rotational agriculture, may have greater success (Akyoo and Lazaro 2007).

### 5.2 Implications for land system science and social-ecological systems

Social-ecological systems, and therefore land systems, are inherently very complex. They commonly exhibit feedbacks between system components and past and future states. Because of this complexity, researchers generally describe phenomena of interest qualitatively, or they break the components of a given causal pathway down into many sub components (Patrick Meyfroidt 2016; B. Turner et al. 2020). Nevertheless, inference from limited time series data is difficult due to issues of equifinality, simultaneous causation, and unobserved complexity(Barrett 2021; Cumming et al. 2020). We echo the argument of Schlüter and others (2019) that agent/individual-based modeling can allow inference about causality in social-ecological systems and emphasize that this approach is especially powerful when combined with empirical data, as presented here, in order to increase validity and interpretability.

We argue here that individual-based models will be of the greatest utility for land system science when they are combined with a standard, systematic framework for comparing synthetic data to real-world observations. Approximate Bayesian computation may fill this niche as it is relatively straightforward and allows for parameter estimation given even very complex generative models. A key advantage of this method is that it allows modelers to explicitly confront equifinality in a given simulation and to some extent a given empirical system. By explicitly quantifying the range of causal processes that are likely to produce an outcome of interest, researchers will greatly increase the applicability of simulation models to real-world policy decisions (Williams et al. 2020).

### 5.3 Limitations and future work

A limitation of any process model is that they assume that researchers know and can accurately represent causal processes in silico. In the case of models like the individual-based simulation presented here, researchers must abstract down to only key phenomena of interest, eliminating much contextual nuance and again, assuming that we know what matters and what does not. This is a big assumption in social-ecological systems considering that emergent and often unexpected phenomena are a defining feature of the field. This limitation is exemplified by the four *shehia* (Fundo, Muambe, Jombwe, and Shamiani) that showed fewer coral rag forest pixels converted to agriculture than expected by the model even under no externally driven forest conversion pressure. We know through our in-person observations and interviews with Community Forests Pemba staff that these four *shehia* have been the focus of considerable tree planting, particularly of *Casuarina spp.* in woodlots. We are likely observing both some confusion between native coral rag vegetation and woodlot vegetation in our satellite observations, as well as a reduction in coral rag deforestation for fuelwood. Approximately 95% of Pemban households rely on cutting fuelwood for daily cooking activities. Hence, the introduction of woodlots, which we do not account for in our simulation, likely reduces the overall rates of coral rag deforestation and conversion to rotational agriculture. Further, Fundo and Shamiani are both islets of Pemba and are both home to luxury resorts. The limited connection between these islets and the main population centers of Pemba likely limits the effect of market forces driving agricultural expansion in these two areas. Also, our anecdotal experience is that the resort operators intentionally limit local development nearby the properties, possibly reducing the rate of agricultural development below what we expect as a result of soil degradation.

Another contextual limitation of our specific model for drawing inference about agricultural expansion in Pemba Island is that we do not account for the long-term processes of soil degradation that lead to complete land abandonment and eventually the recovery of coral rag vegetation. While the two year farm and three year fallow cycle is standard in the coral rag geological areas of Pemba, some agricultural areas are completely abandoned when crop yields are consistently low even after a fallow period. Additionally, the agricultural units themselves are independent 20m pixels which is considerably smaller than many agricultural operations. Clumping these pixels to better match realistic farm sizes may produce different and more accurate inference than presented here.

Lastly, our model does not allow for heterogeneity or evolution in human behavior. All agricultural pixels follow the same scheduling process. While this scheduling process is standard in the study *shehia*, an interesting exercise would be to allow for the diffusion of regenerative agricultural practices across farm units to examine direct feedbacks between environmental and cultural change.

## 6 Conclusion

Computer simulations are critical to theoretical development in land system science as they allow us to formally define and scrutinize hypothesized mechanisms driving phenomena of interest. When we develop competing plausible mechanisms however, it can be difficult to identify the contribution of each hypothesized mechanism in the real world. Recent advances in ABC, primarily in population genetics, but also cultural evolution, have provided a structured process to begin to overcome this challenge in other fields (Hartig et al. 2011; Kandler and Powell 2018). Until now however, ABC has yet to be applied to land system science. In this paper we show how ABC can be used to better leverage the wealth of available satellite data in combination with individual-based models of land system change in order to assess the importance of competing mechanisms.

In particular, we develop an individual-based simulation of agricultural expansion in Pemba, Tanzania under two different mechanisms: soil degradation and external forces such as population growth and increasing market integration. We use ABC to systematically compare runs from this model with observed land cover change in 19 *shehia* in Pemba from 2018 to 2021. This process allows us to estimate the likelihood that various rates of externally driven agricultural expansion are responsible for the observed land cover change in each *shehia*. Importantly, this process also allows us to directly estimate the range of externally driven expansion rates that could have also reasonably resulted in the observed data, or the degree to which the system is equifinal.

## 7 Acknowledgements

We thank the entire Community Forests Pemba staff for continued support on research in Pemba. We also thank the Hazards and Climate Resilience Institute at Boise State University for funding this research. We thank Dr. Monique Borgerhoff Mulder and Dr. Tim Caro their help in conceptualizing the project and with all things related to fieldwork in Pemba. Lastly, we thank the Max Planck Institute for Evolutionary Anthropology, department of Human Behavior, Ecology and Culture for providing the computing resources necessary for this project and Dr. Anne Kandler for her support with the ABC and for providing an initial review of the paper.

## 8 Open data and software

All data, R code, and GEE code used in this project are available at this github: https://github.com/matthewclark1223/CPR_ABM/tree/master/HCRI_Grant and this GEE link: https://code.earthengine.google.com/3774c9546268839fcbf43176e4d2eb46. An open source web app that displays the runs from the agent-based model for specific *shehia* can be found here: https://matthewclark.shinyapps.io/LandUsePredictionsApp/.

## Supplemental

All supplemental materials can be found in the GoogleSheet here: https://docs.google.com/spreadsheets/d/18jGNxQl8uVK2m-jHzR6Pq4H-OmEXWMJFTSswoklx1-8/edit?usp=sharing.

## References

10 Ahimbisibwe, Vianny, Jürgen Groeneveld, Melvin Lippe, Susan Balaba Tumwebaze, Eckhard Auch, and Uta Berger. 2021. “Understanding Smallholder Farmer Decision Making in Forest Land Restoration Using Agent-Based Modeling.” Socio-Environmental Systems Modelling 3 (November): 18036–36. https://doi.org/10.18174/sesmo.2021a18036.

Akyoo, Adam, and Evelyne Lazaro. 2007. “The Spice Industry in Tanzania: General Profile, Supply Chain Structure, and Food Standards Compliance Issues.” Working Paper 2007:8. DIIS Working Paper. https://www.econstor.eu/handle/10419/84561.

An, Li, Volker Grimm, Abigail Sullivan, B. L. Turner II, Nicolas Malleson, Alison Heppenstall, Christian Vincenot, et al. 2021. “Challenges, Tasks, and Opportunities in Modeling Agent-Based Complex Systems.” Ecological Modelling 457 (October): 109685. https://doi.org/10.1016/j.ecolmodel.2021.109685.

Banks, Sarah, Lori White, Amir Behnamian, Zhaohua Chen, Benoit Montpetit, Brian Brisco, Jon Pasher, and Jason Duffe. 2019. “Wetland Classification with Multi-Angle/Temporal SAR Using Random Forests.” Remote Sensing 11 (6): 670. https://doi.org/10.3390/rs11060670.

Barrett, Christopher B. 2021. “On Design-Based Empirical Research and Its Interpretation and Ethics in Sustainability Science.” Proceedings of the National Academy of Sciences 118 (29). https://doi.org/10.1073/pnas.2023343118.

Bastos Lima, Mairon G, U Martin Persson, and Patrick Meyfroidt. 2019. “Leakage and Boosting Effects in Environmental Governance: A Framework for Analysis.” Environmental Research Letters 14 (10): 105006. https://doi.org/10.1088/1748-9326/ab4551.

Beaumont, Mark A, Wenyang Zhang, and David J Balding. 2002. “Approximate Bayesian Computation in Population Genetics.” Genetics 162 (4): 2025–35. https://doi.org/10.1093/genetics/162.4.2025.

Belgiu, Mariana, and Lucian Drăgut. 2016. “Random Forest in Remote Sensing: A Review of Applications and Future Directions.” ISPRS Journal of Photogrammetry and Remote Sensing 114 (April): 24–31. https://doi.org/10.1016/j.isprsjprs.2016.01.011.

Berkes, Fikret, Carl Folke, and Johan Colding. 2000. Linking Social and Ecological Systems: Management Practices and Social Mechanisms for Building Resilience. Cambridge University Press.

Biazin, Birhanu, Geert Sterk, Melesse Temesgen, Abdu Abdulkedir, and Leo Stroosnijder. 2012. “Rainwater Harvesting and Management in Rainfed Agricultural Systems in SubSaharan Africa-a Review.” Physics and Chemistry of the Earth, Parts A/B/C, Recent advances in water resources management, 47-48 (January): 139–51. https://doi.org/10.1016/j.pce.2011.08.015.

Boult, Victoria L., Tristan Quaife, Vicki Fishlock, Cynthia J. Moss, Phyllis C. Lee, and Richard M. Sibly. 2018. “Individual-Based Modelling of Elephant Population Dynamics Using Remote Sensing to Estimate Food Availability.” Ecological Modelling 387 (November): 187–95. https://doi.org/10.1016/j.ecolmodel.2018.09.010.

Breiman, Leo. 2001. “Random Forests.” Machine Learning 45 (1): 5–32. https://doi.org/10.1023/A:1010933404324.

Burgess, N. D., and G. P. Clarke. 2000. “Coastal Forests of Eastern Africa.” Coastal Forests of Eastern Africa. https://www.cabdirect.org/cabdirect/abstract/20013178223.

Burrows, John Eric, Sandie Burrows, Ernst Schmidt, Mervyn Lotter, and Edward O Wilson. 2018. Trees and Shrubs Mozambique. Print Matters Heritage Cape Town.

Casetti, Emilio, and Howard L. Gauthier. 1977. “A Formalization and Test of the “Hollow Frontier” Hypothesis.” Economic Geography 53 (1): 70–78. https://doi.org/10.2307/142807.

Cipriotti, Pablo A., Martín R. Aguiar, Thorsten Wiegand, and José M. Paruelo. 2012. “Understanding the Long-Term Spatial Dynamics of a Semiarid Grass-Shrub Steppe Through Inverse Parameterization for Simulation Models.” Oikos 121 (6): 848–61. https://doi.org/10.1111/j.1600-0706.2012.20317.x.

Cumming, G. S., G. Epstein, J. M. Anderies, C. I. Apetrei, J. Baggio, Ö. Bodin, S. Chawla, et al. 2020. “Advancing Understanding of Natural Resource Governance: A Post-Ostrom Research Agenda.” Current Opinion in Environmental Sustainability, Resilience and complexity:frameworks and models to capture social-ecological interactions, 44 (June): 26–34. https://doi.org/10.1016/j.cosust.2020.02.005.

DeAngelis, Donald L., and Volker Grimm. 2014. “Individual-Based Models in Ecology After Four Decades.” F1000Prime Reports 6 (June): 39. https://doi.org/10.12703/P6-39.

DeFries, Ruth, and J. R. G. Townshend. 1994. “NDVI-Derived Land Cover Classifications at a Global Scale.” International Journal of Remote Sensing 15 (17): 3567–86. https://doi.org/10.1080/01431169408954345.

Epstein, Joshua M. 2008. “Why Model?” Text.Article. October 31, 2008. https://www.jasss.org/11/4/12.html.

Farr, Tom G., Paul A. Rosen, Edward Caro, Robert Crippen, Riley Duren, Scott Hensley, Michael Kobrick, et al. 2007. “The Shuttle Radar Topography Mission.” Reviews of Geophysics 45 (2). https://doi.org/10.1029/2005RG000183.

Folke, Carl. 2007. “Social–Ecological Systems and Adaptive Governance of the Commons.” Ecological Research 22 (1): 14–15. https://doi.org/10.1007/s11284-006-0074-0.

Gallagher, Cara A., Magda Chudzinska, Angela Larsen-Gray, Christopher J. Pollock, Sarah N. Sells, Patrick J. C. White, and Uta Berger. 2021. “From Theory to Practice in Pattern-Oriented Modelling: Identifying and Using Empirical Patterns in Predictive Models.” Biological Reviews 96 (5): 1868–88. https://doi.org/10.1111/brv.12729.

Gao, Bo-cai. 1996. “NDWI—a Normalized Difference Water Index for Remote Sensing of Vegetation Liquid Water from Space.” Remote Sensing of Environment 58 (3): 257–66. https://doi.org/10.1016/S0034-4257(96)00067-3.

Garrity, Dennis Philip, Festus K. Akinnifesi, Oluyede C. Ajayi, Sileshi G. Weldesemayat, Jeremias G. Mowo, Antoine Kalinganire, Mahamane Larwanou, and Jules Bayala. 2010. “Evergreen Agriculture: A Robust Approach to Sustainable Food Security in Africa.” Food Security 2 (3): 197–214. https://doi.org/10.1007/s12571-010-0070-7.

Gorelick, Noel, Matt Hancher, Mike Dixon, Simon Ilyushchenko, David Thau, and Rebecca Moore. 2017. “Google Earth Engine: Planetary-Scale Geospatial Analysis for Everyone.” Remote Sensing of Environment, Big remotely sensed data: Tools, applications and experiences, 202 (December): 18–27. https://doi.org/10.1016/j.rse.2017.06.031.

Grimm, Volker. 1999. “Ten Years of Individual-Based Modelling in Ecology: What Have We Learned and What Could We Learn in the Future?” Ecological Modelling 115 (2): 129–48. https://doi.org/10.1016/S0304-3800(98)00188-4.

Grimm, Volker, and Steven F. Railsback. 2013. Individual-Based Modeling and Ecology. Princeton University Press. https://doi.org/10.1515/9781400850624.

Grimm, Volker, Eloy Revilla, Uta Berger, Florian Jeltsch, Wolf M. Mooij, Steven F. Railsback, Hans-Hermann Thulke, Jacob Weiner, Thorsten Wiegand, and Donald L. DeAngelis. 2005. “Pattern-Oriented Modeling of Agent-Based Complex Systems: Lessons from Ecology.” Science 310 (5750): 987–91. https://doi.org/10.1126/science.1116681.

Hartig, Florian, Justin M. Calabrese, Björn Reineking, Thorsten Wiegand, and Andreas Huth. 2011. “Statistical Inference for Stochastic Simulation Models – Theory and Application.” Ecology Letters 14 (8): 816–27. https://doi.org/10.1111/j.1461-0248.2011.01640.x.

Kaimowitz, David, and Arild Angelsen. 1998. Economic Models of Tropical Deforestation: A Review. CIFOR.

Kandler, Anne, and Adam Powell. 2018. “Generative Inference for Cultural Evolution.” Philosophical Transactions of the Royal Society B: Biological Sciences 373 (1743): 20170056. https://doi.org/10.1098/rstb.2017.0056.

Kaplan, Gregoriy, Lior Fine, Victor Lukyanov, V. S. Manivasagam, Josef Tanny, and Offer Rozenstein. 2021. “Normalizing the Local Incidence Angle in Sentinel-1 Imagery to Improve Leaf Area Index, Vegetation Height, and Crop Coefficient Estimations.” Land 10 (7): 680. https://doi.org/10.3390/land10070680.

Kingdon, Jonathan. 1988. East African Mammals: An Atlas of Evolution in Africa, Volume 3, Part a: Carnivores. University of Chicago Press.

Kosmala, Margaret, Philip Miller, Sam Ferreira, Paul Funston, Dewald Keet, and Craig Packer. 2016. “Estimating Wildlife Disease Dynamics in Complex Systems Using an Approximate Bayesian Computation Framework.” Ecological Applications 26 (1): 295–308. https://doi.org/10.1890/14-1808.

Kramer-Schadt, Stephanie, Eloy Revilla, Thorsten Wiegand, and Urs Breitenmoser. 2004. “Fragmented Landscapes, Road Mortality and Patch Connectivity: Modelling Influences on the Dispersal of Eurasian Lynx.” Journal of Applied Ecology 41 (4): 711–23. https://doi.org/10.1111/j.0021-8901.2004.00933.x.

Lambin, Eric F., and Patrick Meyfroidt. 2010. “Land Use Transitions: Socio-Ecological Feedback Versus Socio-Economic Change.” Land Use Policy, Forest transitions, 27 (2): 108–18. https://doi.org/10.1016/j.landusepol.2009.09.003.

Le, Quang Bao, Roman Seidl, and Roland W. Scholz. 2012. “Feedback Loops and Types of Adaptation in the Modelling of Land-Use Decisions in an Agent-Based Simulation.” Environmental Modelling & Software 27-28 (January): 83–96. https://doi.org/10.1016/j.envsoft.2011.09.002.

Levin, Simon, Tasos Xepapadeas, Anne-Sophie Crépin, Jon Norberg, Aart de Zeeuw, Carl Folke, Terry Hughes, et al. 2013. “Social-Ecological Systems as Complex Adaptive Systems: Modeling and Policy Implications.” Environment and Development Economics 18 (2): 111–32. https://doi.org/10.1017/S1355770X12000460.

Liu, Jianguo, Thomas Dietz, Stephen R. Carpenter, Marina Alberti, Carl Folke, Emilio Moran, Alice N. Pell, et al. 2007. “Complexity of Coupled Human and Natural Systems.” Science 317 (5844): 1513–16. https://doi.org/10.1126/science.1144004.

Martínez, Isabel, Thorsten Wiegand, J. Julio Camarero, Enric Batllori, and Emilia Gutiérrez. 2011. “Disentangling the Formation of Contrasting Tree-Line Physiognomies Combining Model Selection and Bayesian Parameterization for Simulation Models.” The American Naturalist 177 (5): E136–52. https://doi.org/10.1086/659623.

Meyfroidt, Patrick. 2016. “Approaches and Terminology for Causal Analysis in Land Systems Science.” Journal of Land Use Science 11 (5): 501–22. https://doi.org/10.1080/1747423X.2015.1117530.

Meyfroidt, P., J. Börner, R. Garrett, T. Gardner, J. Godar, K. Kis-Katos, B. S. Soares-Filho, and S. Wunder. 2020. “Focus on Leakage and Spillovers: Informing Land-Use Governance in a Tele-Coupled World.” Environmental Research Letters 15 (9): 090202. https://doi.org/10.1088/1748-9326/ab7397.

Meyfroidt, P., R. Roy Chowdhury, A. de Bremond, E. C. Ellis, K.-H. Erb, T. Filatova, R. D. Garrett, et al. 2018. “Middle-Range Theories of Land System Change.” Global Environmental Change 53 (November): 52–67. https://doi.org/10.1016/j.gloenvcha.2018.08.006.

Mondal, Pinki, Xue Liu, Temilola E. Fatoyinbo, and David Lagomasino. 2019. “Evaluating Combinations of Sentinel-2 Data and Machine-Learning Algorithms for Mangrove Mapping in West Africa.” Remote Sensing 11 (24): 2928. https://doi.org/10.3390/rs11242928.

Olofsson, Pontus, Giles M. Foody, Stephen V. Stehman, and Curtis E. Woodcock. 2013. “Making Better Use of Accuracy Data in Land Change Studies: Estimating Accuracy and Area and Quantifying Uncertainty Using Stratified Estimation.” Remote Sensing of Environment 129 (February): 122–31. https://doi.org/10.1016/j.rse.2012.10.031.

Ren, Yanjiao, Yihe Lü, Alexis Comber, Bojie Fu, Paul Harris, and Lianhai Wu. 2019. “Spatially Explicit Simulation of Land Use/Land Cover Changes: Current Coverage and Future Prospects.” Earth-Science Reviews 190 (March): 398–415. https://doi.org/10.1016/j.earscirev.2019.01.001.

Rossmanith, Eva, Niels Blaum, Volker Grimm, and Florian Jeltsch. 2007. “Pattern-Oriented Modelling for Estimating Unknown Pre-Breeding Survival Rates: The Case of the Lesser Spotted Woodpecker (Picoides Minor).” Biological Conservation 135 (4): 555–64. https://doi.org/10.1016/j.biocon.2006.11.002.

Schlüter, Maja, Kirill Orach, Emilie Lindkvist, Romina Martin, Nanda Wijermans, Örjan Bodin, and Wiebren J. Boonstra. 2019. “Toward a Methodology for Explaining and Theorizing about Social-Ecological Phenomena.” Current Opinion in Environmental Sustainability 39 (August): 44–53. https://doi.org/10.1016/j.cosust.2019.06.011.

Schuster, Christian, Michael Förster, and Birgit Kleinschmit. 2012. “Testing the Red Edge Channel for Improving Land-Use Classifications Based on High-Resolution Multi-Spectral Satellite Data.” International Journal of Remote Sensing 33 (17): 5583–99. https://doi.org/10.1080/01431161.2012.666812.

Scranton, Katherine, Jonas Knape, and Perry de Valpine. 2014. “An Approximate Bayesian Computation Approach to Parameter Estimation in a Stochastic Stage-Structured Population Model.” Ecology 95 (5): 1418–28. https://doi.org/10.1890/13-1065.1.

Steffen, Will, Angelina Sanderson, Peter Tyson, Jill Jäger, Pamela Matson, Berrien Moore Iii, Frank Oldfield, et al. 2006. “Global Change and the Earth System: A Planet Under Pressure.” Global Change and the Earth System, 44.

Stehman, Stephen V., and Giles M. Foody. 2019. “Key Issues in Rigorous Accuracy Assessment of Land Cover Products.” Remote Sensing of Environment 231 (September): 111199. https://doi.org/10.1016/j.rse.2019.05.018.

Stockley, Gordon Murray. 1928. Report on the Geology of the Zanzibar Protectorate. Government of Zanzibar.

Sunnåker, Mikael, Alberto Giovanni Busetto, Elina Numminen, Jukka Corander, Matthieu Foll, and Christophe Dessimoz. 2013. “Approximate Bayesian Computation.” PLOS Computational Biology 9 (1): e1002803. https://doi.org/10.1371/journal.pcbi.1002803.

Swanack, Todd M., William E. Grant, and Michael R. J. Forstner. 2009. “Projecting Population Trends of Endangered Amphibian Species in the Face of Uncertainty: A Pattern-Oriented Approach.” Ecological Modelling 220 (2): 148–59. https://doi.org/10.1016/j.ecolmodel.2008.09.006.

Troost, Christian, Andrew Reid Bell, Hedwig van Delden, Robert Huber, Tatiana Filatova, Quang Bao Le, Melvin Lippe, et al. 2022. “How to Keep It Adequate: A Validation Protocol for Agent-Based Simulation.” SSRN Electronic Journal. https://doi.org/10.2139/ssrn.4161475.

Turner, B. L., Eric F. Lambin, and Anette Reenberg. 2007. “The Emergence of Land Change Science for Global Environmental Change and Sustainability.” Proceedings of the National Academy of Sciences 104 (52): 20666–71. https://doi.org/10.1073/pnas.0704119104.

Turner, B. L., Eric F. Lambin, and Peter H. Verburg. 2021. “From Land-Use/Land-Cover to Land System Science.” Ambio 50 (7): 1291–94. https://doi.org/10.1007/s13280-021-01510-4.

Turner, BL, Patrick Meyfroidt, Tobias Kuemmerle, Daniel Müller, and Rinku Roy Chowdhury. 2020. “Framing the Search for a Theory of Land Use.” Journal of Land Use Science 15 (4): 489–508. https://doi.org/10.1080/1747423X.2020.1811792.

Vaart, Elske van der, Mark A. Beaumont, Alice S. A. Johnston, and Richard M. Sibly. 2015. “Calibration and Evaluation of Individual-Based Models Using Approximate Bayesian Computation.” Ecological Modelling 312 (September): 182–90. https://doi.org/10.1016/j.ecolmodel.2015.05.020.

Vaart, Elske van der, Alice S. A. Johnston, and Richard M. Sibly. 2016. “Predicting How Many Animals Will Be Where: How to Build, Calibrate and Evaluate Individual-Based Models.” Ecological Modelling, Next generation ecological modelling, concepts, and theory: Structural realism, emergence, and predictions, 326 (April): 113–23. https://doi.org/10.1016/j.ecolmodel.2015.08.012.

Verburg, Peter H. 2006. “Simulating Feedbacks in Land Use and Land Cover Change Models.” Landscape Ecology 21 (8): 1171–83. https://doi.org/10.1007/s10980-006-0029-4.

Wiegand, Thorsten, Florian Jeltsch, Ilkka Hanski, and Volker Grimm. 2003. “Using Pattern-Oriented Modeling for Revealing Hidden Information: A Key for Reconciling Ecological Theory and Application.” Oikos 100 (2): 209–22. https://doi.org/10.1034/j.1600-0706.2003.12027.x.

Wild, Robert, Moses Egaru, Mark Ellis-Jones, Barbara Nakangu Bugembe, Ahmed Mohamed, Obadiah Ngigi, Gertrude Ogwok, Jules Roberts, and Sophie Kutegeka. 2020. “Using Inclusive Finance to Significantly Scale Climate Change Adaptation.” In African Handbook of Climate Change Adaptation, edited by Walter Leal Filho, Nicholas Oguge, Desalegn Ayal, Lydia Adelake, and Izael da Silva, 1–26. Cham: Springer International Publishing. https://doi.org/10.1007/978-3-030-42091-8_127-1.

Williams, T. G., S. D. Guikema, D. G. Brown, and A. Agrawal. 2020. “Assessing Model Equifinality for Robust Policy Analysis in Complex Socio-Environmental Systems.” Environmental Modelling & Software 134 (December): 104831. https://doi.org/10.1016/j.envsoft.2020.104831.

Xu, Hanqiu. 2006. “Modification of Normalised Difference Water Index (NDWI) to Enhance Open Water Features in Remotely Sensed Imagery.” International Journal of Remote Sensing 27 (14): 3025–33. https://doi.org/10.1080/01431160600589179.

